# Recruitment of *GRASSY TILLERS1 (GT1)* and *RAMOSA3 (RA3)* into distinct genetic networks in the evolution of grass morphology

**DOI:** 10.1101/2025.08.22.670905

**Authors:** Amber E. de Neve, Olivia A. Kelly, Terice Kelly, Samuel Leiboff, Madelaine E. Bartlett

**Affiliations:** Department of Biology, University of Massachusetts; Amherst, MA, USA; Oregon State University; Corvallis, Oregon, USA; Sainsbury Laboratory Cambridge University (SLCU); Cambridge, UK

## Abstract

Changes in form are driven by the differential, context-dependent regulation of pleiotropic genes. How genetic pleiotropy itself emerges, however, remains unclear. The maize genes *GRASSY TILLERS1* (*GT1*) and *RAMOSA3* (*RA3*) are required for axillary meristem suppression, a deeply conserved trait across angiosperms, and for floral organ suppression, a trait which evolved within the grass family. To determine how these pleiotropic functions are regulated, we first established a high-throughput method for quantitative phenotyping of grass flowers. Using this method, we show that distinct environmental mechanisms regulate axillary meristem versus floral organ suppression. In line with these differences, we find upstream regulation of *GT1* and *RA3* has diverged, consistent with their redeployment in flowers. Our results show that, rather than wholesale adoption of genetic networks, developmental genes can retain ancient functions *and* be recruited into other programs in the evolution of form, thereby increasing genetic pleiotropy.

## Main Text

Rather than generating new proteins through coding sequence changes, morphological evolution often arises from changes in when and where conserved proteins are expressed (King & Wilson, 1975; Carroll, 2005). Mutations that impact gene expression instead of protein structure can generate evolutionary novelty while minimizing harmful side effects, thus reducing the risk of negative mutational pleiotropy from drastic changes to protein coding sequences (Prud’homme *et* al., 2007). Pleiotropy, the influence of a single locus on multiple traits, exists in two forms: mutational pleiotropy, where single base pair substitutions, insertions, or deletions affect multiple genetic functions; and genetic pleiotropy, in which null mutations abolish all functions of a gene (Stearns, 2010; Zhang, 2023). While both mutational and genetic pleiotropy can be positive (or adaptive) for evolution (Foster et al., 2004; Kostyun et al., 2019; Hämälä et al., 2020), negative (or antagonistic) pleiotropy occurs when evolution is constrained due to phenotypic effects that both increase and decrease fitness (Zhang, 2023). As the number and complexity of traits influenced by pleiotropy increase, beneficial mutations become less likely, as positive effects get diluted by unwanted negative side effects on other traits (Orr, 2000; Welch & Waxman, 2003). This layered complexity often obscures the origins of pleiotropy, which remain difficult to pinpoint.

Pleiotropy may emerge from the same genetic modules being redeployed to shape morphology. For example, parts of the genetic program regulating lateral branch initiation in maize are recruited to promote the growth of the leaf ligule, a specialized hinge structure at the sheath-blade boundary (Johnston et al., 2014; Bertolini et al., 2025). While in this case module redeployment promotes growth, in other contexts modules can be co-opted to suppress it (Klein *et al*., 2022).

In maize, two pleiotropic genes involved in growth suppression are *GRASSY TILLERS1* (*GT1*) and *RAMOSA3* (*RA3*) (Satoh-Nagasawa et al., 2006; Whipple et al., 2011; Klein et al., 2022). *GT1* controls selective pistil suppression in maize tassel flowers, a process that is essential to the development of unisexual staminate flowers (Whipple et al., 2011). Unisexual flowers in the maize tassel initiate both stamen and pistil primordia, then pistil primordia undergo cell death leaving behind only the stamens (Calderon-Urrea & Dellaporta, 1999). Besides its role in flower development, *GT1* is also an important regulator of tillering; *gt1* mutants have pistils in tassel florets, as well as additional tiller branches and husk leaves (Whipple et al., 2011). *RA3* also affects both branching and floral development and encodes a trehalose-6-phosphate (Tre6P) phosphatase (Satoh-Nagasawa et al., 2006; Klein et al., 2022). Notably, double *gt1;ra3* mutants have increased tiller branching compared to *gt1* alone, and fully derepressed pistils in both tassels and ears (Klein et al., 2022). Both GT1 and Tre6P phosphatases like RA3 have ancient roles in regulating tiller suppression. *GT1* homologs suppress tiller branching in brachypodium, rice, and arabidopsis (González-Grandío et al., 2017; Kumar et al., 2021; Gallagher et al., 2023; Sánchez-Gerschon *et* al., 2024). Similarly, Tre6P phosphatases like RA3 regulate branching across species: increased Tre6P phosphatase expression decreases branching in the arabidopsis vasculature (Fichtner et al., 2021), and Tre6P influences branch bud outgrowth initiation in garden pea (Fichtner et al., 2017). Together, *GT1* and *RA3* demonstrate how conserved modules can shape plant form, through their effects on both floral development and tillering.

The evolutionary history of pistil suppression versus tiller suppression is known. Growth suppression of lateral vegetative buds (including grass tillers) is a conserved trait in the angiosperms (Takeda et al., 2003; Aguilar-Martínez et al., 2007; Martín-Trillo et al., 2011; Dixon et al., 2018), while growth suppression of floral organs is derived. Programmed cell death of pistils likely emerged in a lineage leading to the Andropogoneae, the tribe to which maize belongs (Le Roux & Kellogg, 1999). Because these two suppression phenotypes occur within the same organism, regulated by the same genes, maize *gt1* and *ra3* mutants represent an ideal system for dissecting the genetics of tiller and pistil suppression.

In tillering, environmental signals, particularly light and sugar availability, play critical roles in regulating branch suppression (Doust, 2007; Barbier et al., 2015b,a; Demotes-Mainard et al., 2016; Khangura et al., 2020; Patil et al., 2022), but environmental effects on pistil suppression are less clear. *TEOSINTE BRANCHED1 (TB1)* integrates environmental signals to suppress tillering (Doebley et al., 1995; Kebrom et al., 2006). TB1 homolog expression is induced by shade in sorghum and arabidopsis (Kebrom et al., 2006; González-Grandío et al., 2013), and *TB1* in pea plants is downregulated when bud growth is activated by sucrose (Mason et al., 2014). Notably,

*GT1* is a downstream target of TB1 in maize, and TB1 binding peaks are located near *RA3*, which is also downregulated in *tb1* mutants (Dong et al., 2019). *GT1* homolog expression in tillers is induced by shade through TB1 homologs in maize and arabidopsis (Whipple et al., 2011; González-Grandío et al., 2017). The TB1-*GT1* interaction is deeply conserved (González-Grandío et al., 2017), as is environmental plasticity of branching (Rameau et al., 2014; Luo et al., 2021). Environmental conditions can also affect pistil suppression (Nuccio et al., 2015; Qian et al., 2025), but whether the same stressors and upstream regulators as in tillering are involved is unknown.

To assess whether TB1 and environmental signals are upstream of pistil suppression as they are in tillers, we developed a sensitive, high-throughput method that can identify subtle morphological changes in pistils. By examining the impact of defoliation and *tb1* mutant combinations on pistil development, we uncover how novel pleiotropies emerge.

### X-ray imaging paired with automated segmentation captures pistil shape data at scale

Measuring maize pistils has been difficult, because each pistil needed to be dissected out of the spikelet (Klein et al., 2022). To address this problem and to investigate the effects of the environment on sporophytic sex determination in maize, we developed a method that combines x-ray imaging and machine learning-based segmentation to measure pistils. We grew maize plants and collected the lowest tassel branch when the anthers emerged from the spikelets and prepared them for x-ray CT (Computed Tomography; Fig. 1a). We chose this collection point 1) to standardize inflorescence maturity, and 2) when the anthers are outside of the spikelet, pistils are clearer on the x-ray image. Because the pistil tissue has high density compared to the surrounding spikelet and branch tissue, the pistils absorb more x-rays and are distinguishable as dark circles in images (Fig. 1b). Instead of reconstructing full 3D volumes, we collected 2D x-ray scans from a reduced number of angles. This approach not only reduced scan time (<5 minutes per tassel branch), but also produced higher-quality images for CNN segmentation than 3D reconstructions, enabling more accurate detection of pistils.

**Figure 1.**
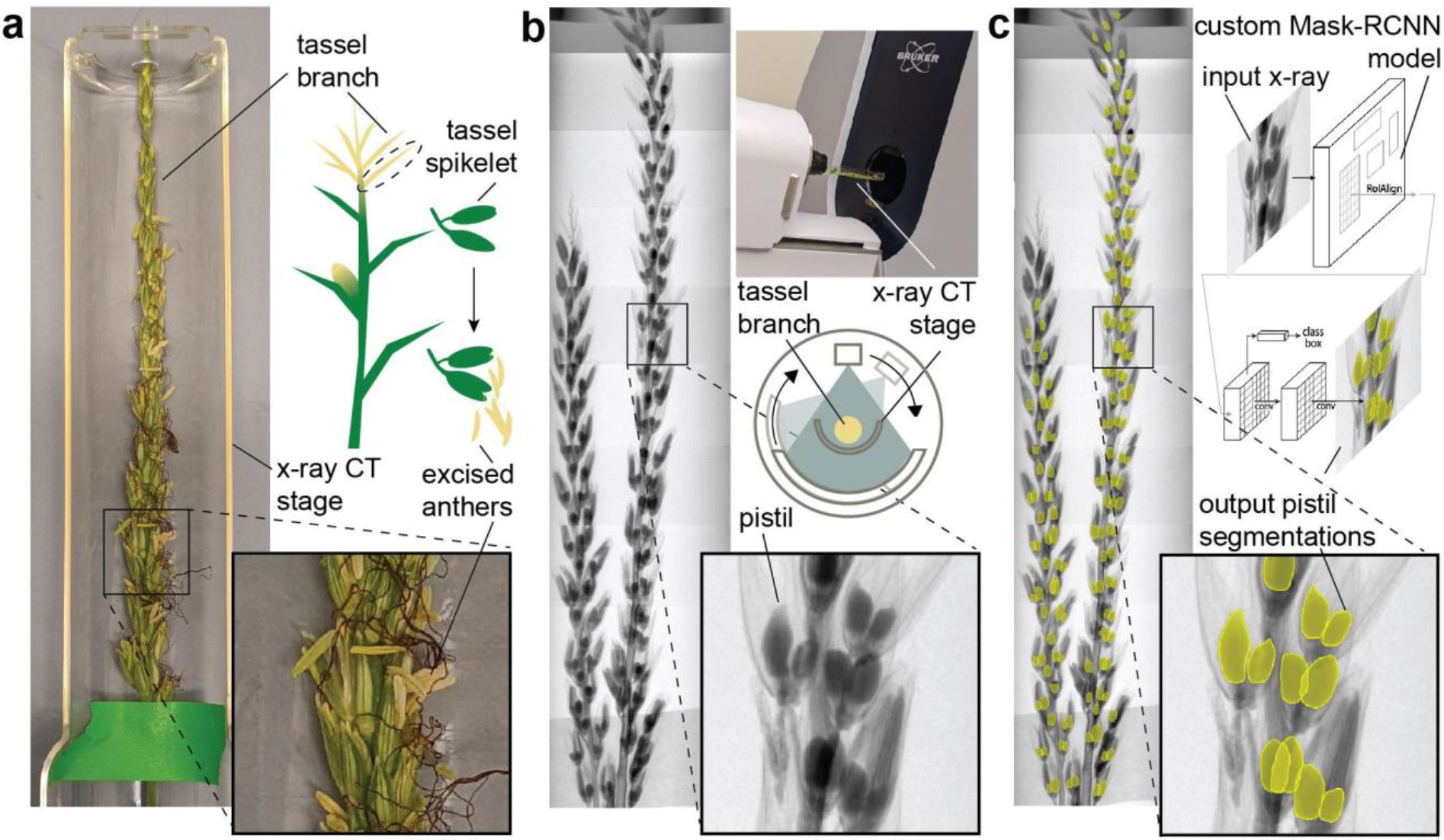
X-ray imaging and automated segmentation quantifies grass floral morphology. a) The bottom tassel branch of a maize plant is collected when anthers are fully emerged from spikelets, and secured to the x-ray CT stage. Ex. *gt1;ra3* maize plant. b) The branch is loaded horizontally into the x-ray CT machine and scans are collected along the length of the sample. Individual scans stitched into whole tassel images. c) Scans are processed by a custom trained Mask-RCNN model to segment pistils (diagram adapted from He et al. 2017), and data from masks are used in subsequent statistical analysis.

Next, we trained a Mask-RCNN (Region-based Convolutional Neural Network) model to automatically measure pistils from the x-ray images (He et al., 2017). Mask-RCNN models can detect objects in an image and generate segmentation masks of those objects (Fig. 1c). To train the model, we first generated a training data set of 664 x-ray images, including maize mutants with defects in tassel pistil repression and samples of *Setaria viridis* (setaria), a grass where pistils are not repressed under normal conditions (Supplementary Table 1). By incorporating setaria, we built an evolutionarily informed model with enhanced applicability across grasses. We validated the performance of the model (Supplementary Fig. 1; Supplementary Tables 2-3), and determined that applying a confidence threshold of 0.82 maximized the precision (fraction of correct predictions out of all predictions) and recall (fraction of correctly identified positives out of all positives) (Rainio et al., 2024) of high overlap predictions (Supplementary Fig. 1). At the 0.82 confidence threshold, the median Intersection over Union (IoU), a measure of overlap between predicted and ground truth segmentations, was 0.83 (IQR = 0.73 - 0.89), indicating high accuracy for most predicted pistil segmentations (Supplementary Fig. 1). The model achieved an average precision (AP) of 0.96 at an IoU threshold of 0.50 (AP_50_), and a mean average precision (mAP) of 0.70 across IoU thresholds from 0.50 to 0.95 in 0.05 increments (Supplementary Fig. 1). Importantly, the model generalized across multiple genotypes, developmental stages, and species, indicating that we built a robust tool for quantifying pistil morphology in grasses.

### Mutant pistils are morphologically varied, and pistil size is affected by spikelet and flower position on the tassel branch

We first used our new method to characterize morphological variability in maize mutants. Floral morphological traits vary along inflorescence axes (Diggle, 1995; Egger & Walbot, 2015).To determine how pistil morphology changed along tassel branches in maize, we assessed the distal-proximal position of pistils on tassel branches. All tassel branch lengths were normalized to 1, and pistil position ranged between 0 at the base - 1 at the tip (Fig. 2a). We quantified pistil presence/absence along branch axes in both *gt1* and *gt1;ra3* mutants. Ectopic derepressed pistils were present along the whole tassel branch for both genotypes, with fewer pistils found in the top 10% of the branches (Fig. 2b). *gt1* mutants often had just one visible derepressed pistil per spikelet compared to *gt1;ra3* mutants which most often had two clearly visible pistils per spikelet (Fig. 2a,b). Along with fewer pistils, *gt1* tassel branches had fewer total spikelets per branch than *gt1;ra3* (Fig. 2b), likely due to a difference in branch length.

**Figure 2.**
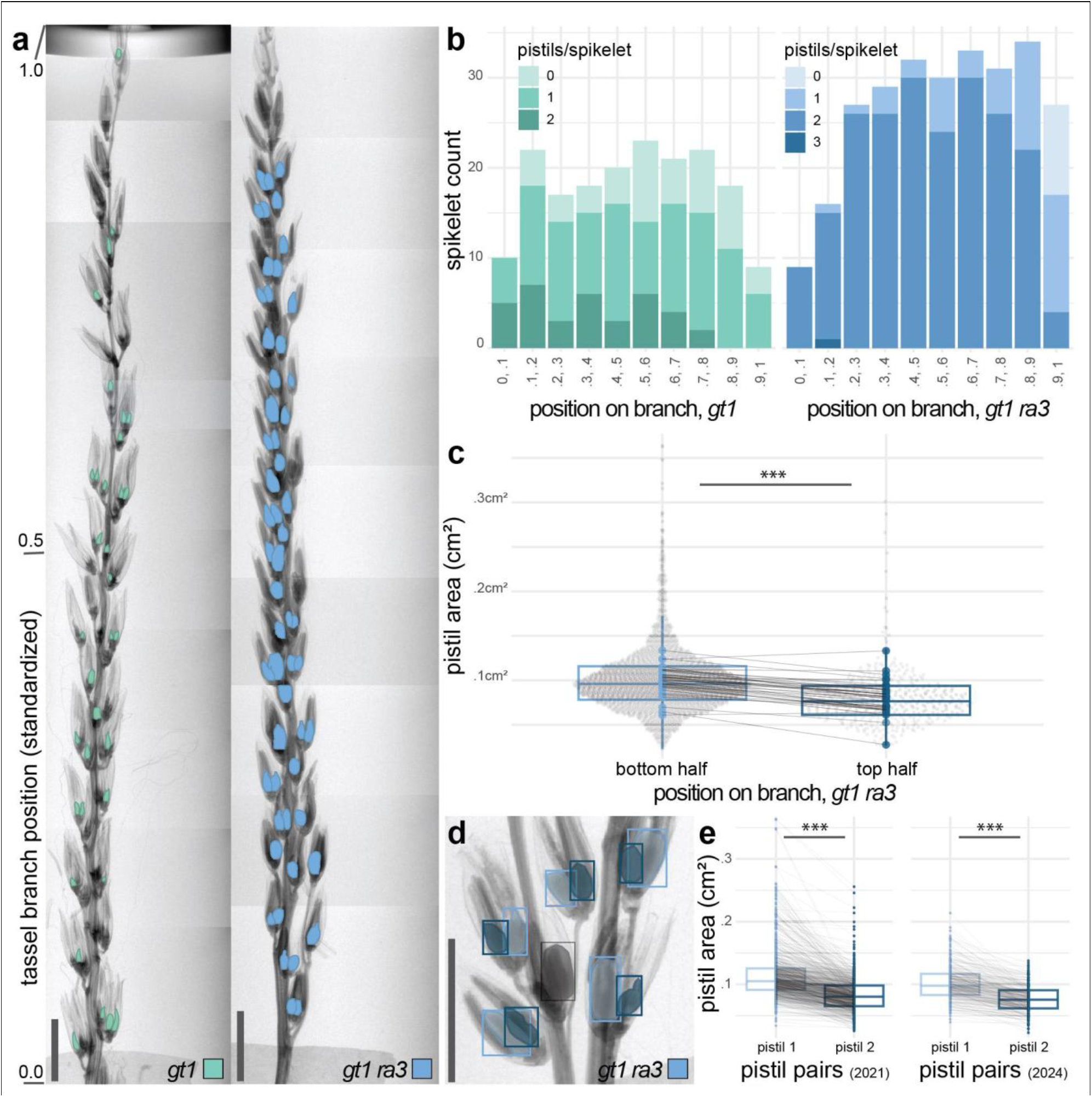
Pistil size changes by branch position. a) Examples of *gt1* and *gt1;ra3* tassel branch x-rays, showing pistil detection masks. The position of pistils on the tassel branch is standardized for each branch, with 0 being the base and 1.0 being the top. Scale bars = 1.9 cm. b) Pistil presence/absence by tassel branch position. Pistils per spikelet for *gt1* and *gt1;ra3* plants. 5 plants measured per genotype (see Methods). c) Pistil area by tassel branch position. *gt1;ra3* tassel branches, bottom halves compared to top halves. Large dots represent the mean pistil area for the branch half, lines connect the same branch. Linear mixed model, pistil area ∼ branch position (bottom/top) * block + (1 | plant). Estimated marginal means, p < .05, ***. d) Example of pistil spikelet pair detections - light blue, larger of pair; dark blue, smaller of pair; black, filtered (not part of a detection pair). e) *gt1;ra3* pistil pair size comparison within spikelets. Lines connect the pistil pairs, split by year. Student’s t-test; p < .05, ***.

Next, using just the *gt1;ra3* dataset, we quantified pistil size along branches. Pistils at the base of tassel branches were larger than those at the top (base pistils = 0.106 cm^2^ ± 0.002; top pistils = 0.084 cm^2^ ± 0.004) (Fig. 2c). This is reminiscent of *Ranunculus, Solanum* and orchid species, where distal flowers are smaller than proximal flowers on branches, and some distal flowers are unable to set fruit (Diggle, 1995; Miller & Diggle, 2003; Diggle & Miller, 2004). Along with positional variability across tassel branches, we also found variation in pistil size within spikelets (Fig. 2d,e). Tassel spikelets have two staminate flowers with suppressed pistils, whereas in *gt1;ra3* plants both pistils develop (Klein et al., 2022). To measure differences at the spikelet level, we isolated all pistil pairs (see Methods). One pistil of the pair was on average ∼25% smaller than the other (pistil 1 = 0.114 cm^2^ ± 0.002; pistil 2 = 0.083 cm^2^ ± 0.002) (Fig. 2d,e). This is consistent with previous data showing maize upper floret anthers are roughly twice the size of the lower floret anthers within a spikelet at any point on the tassel (Egger & Walbot, 2015). The upper flower is larger and develops about a day ahead of the lower flower, and genes are differentially expressed between the two floret types (Yang et al., 2022), indicating upper and lower flowers may have developmental differences.

Because our method captures the outlines of pistils, we could also measure how shape differs across genotypes. We assayed various shape metrics, including elliptical Fourier shape descriptors (Supplementary Fig. 2b), which revealed phenotypes in *gt1;ra3* and *tb1;gt1;ra3*, including rare instances of triple pistils within spikelets and pistils with abnormal morphology (Supplementary Fig. 2c). Upon closer inspection of those flowers with abnormal pistil morphology, likely nucellus tissue and an additional membrane (possibly an ovule integument) were overgrown and protruding out of the outer carpels, which were not fully fused (Supplementary Fig. 2d). This means the chalaza, the ovule-producing meristem (Rudall, 1997; Gross-Hardt et al., 2002), is likely affected in these mutants.

### Defoliation affects pistil and branch suppression differently

We noticed a small but significant difference in pistil size in *gt1;ra3* double mutants between years (2021 *gt1;ra3* = 0.101 cm^2^ ± 0.002; 2024 *gt1;ra3* = 0.091 cm^2^ ± 0.002), suggesting that pistil development is environmentally sensitive (Fig 2c,e), consistent with findings in the ear, the axillary inflorescence of maize (Setter et al., 2001; Gustin et al., 2018). Given that tillering is so sensitive to the environment (Tetio-Kagho & Gardner, 1988; Doebley et al., 1995), and given that this environmental sensitivity feeds through *TB1 (Whipple et al*., *2011; González-Grandío et al*., *2017)*, we wondered if environmental regulation of tiller and pistil development was similar. To assess this, we defoliated *gt1;ra3* plants during development, a treatment known to decrease tillering in maize, likely because of sugar deprivation (Kebrom et al., 2010; Kebrom & Mullet, 2015).

If the environmental sensitivity of pistil suppression and tillering acted through the same pathways, we expect reduced pistil growth in tassels alongside reduced branching. While defoliation decreased tillering as expected (no treatment = 1.08 ± 0.152; defoliated = 0.673 ± 0.128; n = 37) (Fig. 3b), and decreased inflorescence branching (no treatment = 7.11 ± 0.62; defoliated = 5.06 ± 0.57; n = 37) (Fig. 3a), pistil size and shape were not significantly different between defoliated and untreated plants (no treatment = 0.100 cm^2^ ± 0.001; defoliated = 0.098 cm^2^ ± 0.002; tassel n = 37; no treatment pistil n = 2892; defoliated pistil n = 1558) (Fig. 3c and Supplementary Fig. 3). Instead, pistil size and shape stayed the same between treatments. This result was remarkably consistent across two experimental replicates, over two field seasons. Thus, pistils in *gt1;ra3* tassels are resilient to sugar deprivation under conditions that decrease the activity of other branching axes. One possibility is that TB1 or other upstream regulators feed only through *GT1* and *RA3* in environmentally mediated pistil suppression, explaining the absence of a pistil phenotype in *gt1;ra3* mutants. Alternatively, pistil and branch suppression may respond differently to defoliation and upstream regulation of suppression has diverged. In either case, these results indicate that suppression is controlled differently in pistils versus branches.

**Figure 3.**
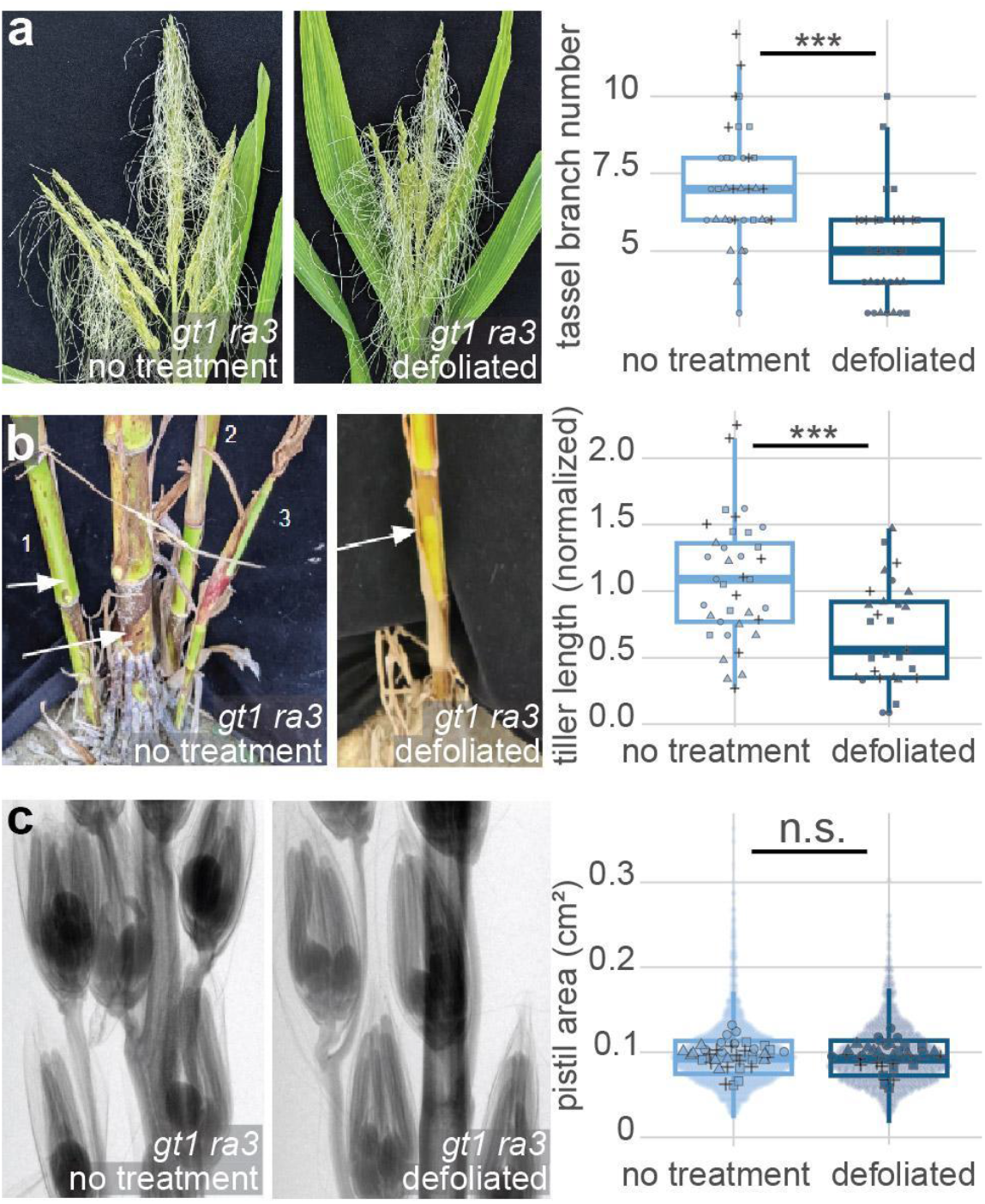
Defoliation affects tassel branch number and tiller length, but not pistil size. a) *gt1;ra3* tassels of untreated and defoliated plants. b) Bases of *gt1;ra3* plants, arrow indicates main stalk, numbers indicate tillers. Tassel branching and tillering significance tested with a 2-way ANOVA assessing block and treatment effects. *** = p-value < .05. c) Pistil x-rays from *gt1;ra3* tassels. Significance tested with a mixed linear model: pistil area ∼ treatment * block + (1 | plant). n.s. = not significant.

### The regulation of *GT1* by TB1 is distinct between tillers and pistils

In addition to sugar, light is a major regulator of tiller suppression, acting upstream of TEOSINTE BRANCHED1 (TB1) to promote tiller outgrowth in shade through differential regulation of its downstream target *GT1* (Tetio-Kagho & Gardner, 1988; Doebley et al., 1995; Kebrom et al., 2006; Whipple et al., 2011; Dong et al., 2017, 2019). The TCP family to which *TB1* belongs also has roles in floral organ suppression (Luo et al., 1996; Yant *et* al., 2015; Jiao et al., 2018), raising the possibility that *TB1* could contribute to pistil suppression, perhaps through a similar TB1-*GT1* pathway. However, pistil suppression in tassels is normal in *tb1* single mutants (Hubbard *et al*., 2002), suggesting that other factors could be compensating for the loss of TB1 in this context. We therefore tested if *tb1* affects suppression in different sensitized mutant backgrounds, including *tb1;ra3* double mutants and the *tb1;gt1;ra3* triple.

We grew B73 (WT), *gt1, ra3, tb1, tb1;ra3* and *tb1;gt1;ra3* mutants in the field and x-rayed tassel branches for analysis (Fig. 4a). *tb1;gt1;ra3* mutants had tassel-like ear inflorescences with silks (Supplementary Fig. 4). After filtering out plants with tassels that had fewer than 5 detected pistils, only *gt1, gt1;ra3* and *tb1;gt1;ra3* plants remained. *gt1* pistils were smaller than both *gt1;ra3* and *tb1;gt1;ra3*; however *gt1;ra3* and *tb1;gt1;ra3* pistil sizes were not significantly different (Fig. 4b). *tb1, ra3* and *tb1;ra3* did not have visible pistils at maturity, as confirmed by scanning electron microscopy (SEM, Fig. 4c). Importantly, all detected pistils in *tb1;ra3* double mutants were individually evaluated and clearly represented false positives. Therefore, unlike in vegetative branching, *tb1* does not affect pistil size in *ra3* or *gt1;ra3* backgrounds. The absence of pistils in *tb1;ra3* mutants indicates that TB1 is unlikely to act upstream of *GT1* or *RA3* in tassel pistil suppression (Fig. 5). In addition, similar pistil sizes between *gt1;ra3* and *tb1;gt1;ra3* mutants indicates that TB1 is not suppressing pistil size through a separate pathway to *GT1* and *RA3*. This is distinct to tillering, where TB1 regulates both *GT1* and *TASSELS REPLACE UPPER EARS1 (Dong et al*., *2017, 2019)*.

**Figure 4.**
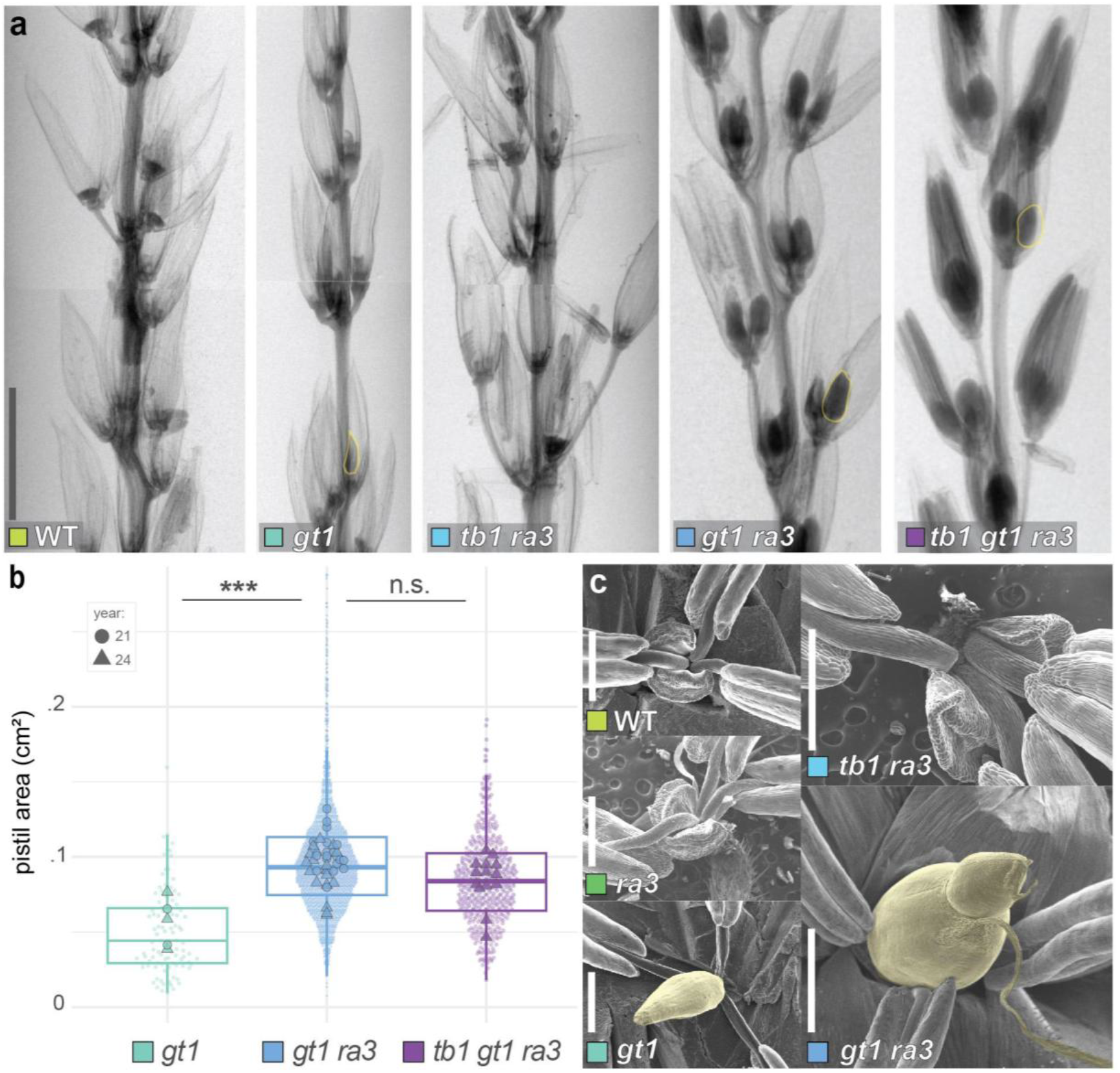
*tb1* does not affect pistil size in *ra3* or *gt1;ra3* backgrounds. a) X-ray tassel sections of WT (B73), *gt1, tb1;ra3, gt1;ra3*, and *tb1;gt1;ra3*. Example pistils are highlighted in yellow. b) Pistil area comparison between *gt1, gt1;ra3* and *tb1;gt1;ra3*. Additional plants were also included in this analysis (B73 - 9, *gt1* - 6, *tb1;ra3* - 2, *tb1* - 13, *ra3* - 2) however they all had less than 5 pistil detections per branch, which was considered the threshold for background noise, and were therefore filtered out. For statistical comparisons, a mixed linear model was used: pistil size ∼ genotype * block + (1 | plant_id) with estimated marginal means. *** = p < .05; n.s. = not significant. c) SEM images of mature tassel florets from various genotypes. Pistils are highlighted with yellow.

**Figure 5.**
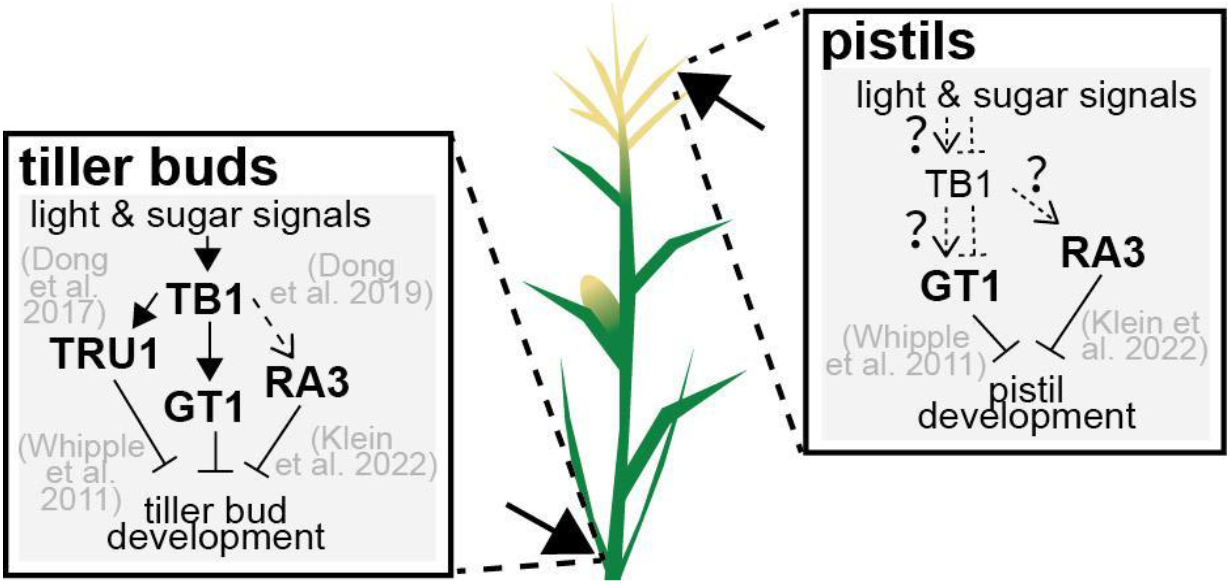
Models for suppression in tiller buds and pistils. Left) Tiller buds are suppressed by *GT1*, which is activated by TB1 in tiller buds in response to light and possibly sugar signals (Whipple et al., 2011; Mason et al., 2014). TB1 promotes the expression of *RA3* (Dong et al., 2019). *RA3* with *GT1* suppresses tiller bud growth (Klein *et al*., 2022). Right) Possible model for pistil suppression.

## Discussion

Conserved developmental modules may have been redeployed in the evolution of pistil suppression, as some genes that suppress tiller growth also suppress pistil growth (Whipple *et al*., 2011). To assess whether an axillary meristem suppression module had been recruited to pistil suppression in the evolution of grass flower morphology, we developed a method to precisely measure the size and shape of pistils in different conditions. Using this method, we found that defoliation, as a proxy for sugar deprivation (Kebrom et al., 2010; Kebrom & Mullet, 2015) does not affect the size or shape of pistils in the tassel, even while it greatly reduces tillering and inflorescence branching. Through mutant comparisons, we determined that *GT1* is likely not downstream of TB1 in pistil suppression. These results indicate that tiller and pistil suppression mechanisms, while controlled by some of the same genes, have distinct upstream regulation.

Scientific progress is often limited by what one can observe; observational gaps can create bottlenecks in understanding plant phenotypes (Atkinson et al., 2019). This especially applies to the study of pistil development in grasses because pistils need to be individually dissected out of tiny spikelets to be measured. Floral development, specifically the formation of pistils vs. stamens in unisexual flowers, is quantitative rather than binary (Lloyd, 1980; Pannell, 2017; Subramaniam & Bartlett, 2023), and the final form a flower takes can depend on several factors such as position (Diggle, 1995; Orr et al., 2001), gene dosage (Moschin *et al*., 2021), and environment (Lan *et al*., 2023). These small quantitative differences in floral development are difficult to measure, especially in grasses. Our approach integrates x-ray imaging with machine learning-based segmentation, making it feasible to measure multiple pistils simultaneously, and extract detailed data regarding their size, shape, and position. Trained across genotypes and species, our model captures features that generalize evolutionarily, extending its utility beyond maize to diverse grasses.

The ability to identify subtle phenotypic changes has broad application in biology, particularly in understanding complex plant traits and their evolution. The omnigenic model proposes that hundreds to thousands of genes influence a trait, but on their own genes may have very weak or no phenotype (Boyle et al., 2017; Mähler et al., 2020). In addition, complex dosage interactions amongst small-effect alleles may generate synergistic phenotypic outcomes (Zebell et al., 2025). Accurately identifying minor phenotypic differences can help detect small-effect alleles, and reveal incremental genetic changes that underpin evolutionary processes and phenotypic diversity. Additionally, finding mutations that have weak effects on a trait can provide new variation for plant breeding programs (Muños et al., 2011; Rodríguez-Leal et al., 2017). While many new technologies can be expensive and inaccessible for labs (Rogers et al., 2024), machine learning models are becoming increasingly accessible (Gehan et al., 2017; Winfree, 2022). This work demonstrates how a fine-tuned computer vision model can be applied to a large image dataset to overcome phenotyping limitations and reveal new things about flower development.

Negative pleiotropy can be circumvented through *cis-*regulatory evolution, as mutations in regulatory elements of pleiotropic genes can restrict changes to a specific tissue and avoid harmful side effects (Carroll, 2008). Our results suggest that this mechanism of avoiding negative pleiotropy might increase the potential for developmental novelty when genes are redeployed across distinct developmental contexts, recruited by different upstream regulators. *GT1* is broadly capable of growth suppression, potentially through jasmonic acid signaling (Klein et al., 2022; Yuan et al., 2025). Supporting this generalizable function, ectopic expression of maize *GT1* in arabidopsis under a floral whorl-specific promoter reduced stamen and petal sizes (Gallagher *et* al., 2023). Likewise, cis-regulatory changes upstream of *GT1* may drive its expression in tassel pistils, where it suppresses growth. Flexible regulatory mechanisms may allow unique roles for genes like *GT1* and *RA3* across developmental contexts, which then manifest as genetic pleiotropy in null mutants. In the grasses in particular, repeated recruitment of growth suppression genes may have led to floral diversification through the suppression of specific floral organs (Komatsuda *et* al., 2007; Klein et al., 2022; Gallagher et al., 2023). In this case, recruitment of a growth suppression module may be a cause - not a consequence - of genetic pleiotropy.

## Materials and Methods

### Plant Material and Growth Conditions

We used the maize genotypes B73, *gt1-P* (Gallagher et al., 2023), *ra3, tb1, tb1 ra3, gt1 ra3*, and *tb1 gt1 ra3* for phenotyping. We used the *gt1* allele *gt1-1* introgressed into B73 (Whipple et al., 2011; Klein et al., 2022). The *ra3* used was *ra3-fea1* (B73 background) (Satoh-Nagasawa et al., 2006). The *tb1* allele used was *tb1-ref* introgressed into B73 (Doebley et al., 1997). Maize was grown at the University of Massachusetts Amherst Crop and Animal Research and Education Farm in South Deerfield, MA (∼42°290N, 72°350W) and setaria plants were grown at the College of Natural Sciences and Education Greenhouse on the UMass Amherst campus under long day conditions (16 h light, 8 h dark) at 28°C. Maize for defoliation experiments was grown at the University of Massachusetts Amherst Crop and Animal Research and Education Farm (∼42°29′N, 72°35′W).

### Defoliation Assay and Phenotyping

Plants were split across different parts of the field in a block design. Every other plant in the row was defoliated by removing half of the most recently exposed leaf starting at 3 weeks after planting and repeating every 3-4 days for 5 weeks. This experiment was repeated for 2 field seasons. Tiller length measurements were standardized to plant height and then averaged per plant.

### X-ray Imaging

X-ray scans were captured using a Bruker Skyscan 1276 uCT at the Animal Imaging Core Facility on the UMass Amherst campus. Inflorescence samples were loaded horizontally into the machine. Samples were x-rayed from a reduced number of angles, and multiple scans were captured along the length of the sample. A single image angle from the top was selected and used in subsequent analyses. Scans were taken at 1008×672 resolution. The zoom level was set at 30μm per pixel for maize, and 40μm per pixel for setaria. All maize x-ray scans were taken at anthesis, and the bottommost tassel branch was used for imaging.

### Pistil segmentation model

To train the model we first generated the training, validation and test data sets (table S1). Maize mutants with defects in tassel pistil repression were grown in either the field (UMass Amherst Crop and Animal Research and Education Farm) or the greenhouse (UMass Amherst College of Natural Sciences and Education Greenhouse). To obtain a range of pistil phenotypes, we grew the inbred line B73 as the negative control, *gt1-P, gt1 ra3* and *tb1 gt1 ra3* mutants. We also included *Setaria viridis* inflorescence examples to make the model more generalizable for pistil detection in other grass species. For these examples, we collected setaria at various growth stages to get a range of pistil sizes for training. Setaria x-rays were collected at 4 developmental stages (28, 31, 33, 42 days after sowing (DAS)). Branches were removed from the setaria inflorescences with tweezers to reduce overlap, and the thinned inflorescence was used for imaging. After obtaining x-ray images for all samples, we annotated pistils by tracing using the Via-annotator program (Dutta & Zisserman, 2019).

We used Roboflow (Dwyer, B., Nelson, J., Solawetz, J., et. al., 2022) to standardize the input image size in the case of different resolutions and to generate synthetic training examples using rotation (+-10 degrees), brightness (+-7%), and blur (up to .5px). The total dataset size was 332 tassel x-rays without synthetic data, and 664 after adding synthetic data. X-ray images were converted from tiff to jpg format, and pistil annotations were checked for accuracy in a Jupyter notebook (available on Github) before training.

Pistils in x-ray images are dense (20+ pistils per image) and overlapping. Because of this, Mask-RCNN (He et al., 2017) was chosen as the base model due to its ability to segment overlapping objects effectively (Rettenberger et al., 2023). Pistil-specific models were trained using Mask R-CNN with a ResNet-101-FPN backbone, pretrained on ImageNet and obtained from the Detectron2 Model Zoo (Yuxin Wu, Alexander Kirillov, Francisco Massa, Wan-Yen Lo, Ross Girshick, 2019). Models were trained using the Detectron2 framework using the default Mask-RCNN hyperparameter configurations. Training was completed on the Unity Cluster located at the Massachusetts Green High Performance Computing Center. During training several metrics were logged every 200 iterations, including total loss, validation loss, bounding box AP and segmentation AP. Model selection was based on the lowest validation loss. Checkpoints were saved every 200 iterations, and the final model was selected at iteration 1399 when validation loss plateaued.

We used stratified 5-fold cross-validation to evaluate model performance (table S2). Data splitting was implemented in Python using StratifiedKFold from scikit-learn, stratified by plant species (maize vs. setaria) and pistil size category (small, medium, large, extra-large). This ensured each fold contained a balanced composition of image types. For each fold, models were trained as described above, and training was stopped at 1399 iterations. Model performance across folds was assessed using bounding box and segmentation precision (AP) and consistency was compared across splits. The final model was evaluated using the hold-out test set with coco evaluation metrics (table S3). Lastly, a confidence threshold of 0.82 was applied to maximize the F1 score at high IoUs (fig. S1).

### Pistil area and shape genotype comparison

Maize plants from all genotypes were grown for pistil area and shape quantification (see Plant Material and Growth Conditions). Tassel x-ray scans were collected as described above. The final pistil segmentation model with the 0.82 confidence threshold applied was used to generate pistil masks from the x-ray scans. First, pistil detections were filtered to remove detections of cut-off pistils at the edges of the image. Duplicate pistils in the regions where successive x-ray scans overlap were also removed. Tassels with less than 5 detections per tassel were filtered from the analysis. Pistil size was calculated from the total pixels in the detection mask. Pistil shape metrics were calculated using Python OpenCV. Fourier shape analysis was done using the program SHAPE (Iwata & Ukai, 2002). To quantify the pistil presence/absence by tassel branch position, model outputs were manually checked against the original x-rays to ensure no pistil detections were missed. To find pistil pairs within spikelets, a custom python script was used to identify instances where only two pistil masks overlapped. Any singletons or overlaps of more than two pistils were discarded from the analysis. Identified pistil pairs were viewed to confirm pairs were correct before downstream analysis. All analyses were performed in Jupyter notebooks or R, and files are available on Github.

### Genotyping Assays

*tb1 ra3* and *tb1 gt1 ra3* plants were genotyped with the following primers:*tb1-ref* forward - GCCTTGGAGTCCCATCAGT, *tb1-ref* reverse - TTCATCGTCACACAGCCAAT; *ra3-fea1* forward CGGGAGGCGGGGAACTAATA, *ra3-fea1* reverse - AGTAGCACTCTGGGAAAGCG. A fragment surrounding the *tb1-ref* or *ra3-fea1* lesion was amplified with Phusion High-Fidelity DNA Polymerase (New England Biolabs) using the supplied HF buffer. Thermal cycling conditions were as follows: initial denaturation at 98 °C for 30 s; 30 cycles of 98 °C for 10 s, primer-specific annealing temperature (*tb1-ref* 63 °C, *ra3-fea1* 64 °C) for 20 s, and 72 °C for 30 s/kb; with a final extension at 72 °C for 5 min. PCR products were purified with the DNA Clean & Concentrator kit (Zymo Research) and submitted for Sanger sequencing.

### Scanning Electron Microscopy (SEM)

SEM was performed with a JEOL JCM-6000 Plus Neoscope Benchtop scanning electron microscope with fresh tissue samples. Late-stage florets were dissected from the tassel and transferred to the SEM. Images were recorded under high vacuum and 5-kV voltage within 15 min of being in the SEM.

## Acknowledgements

We thank D. Chitwood, C. Whipple, E. Kellogg, and A. Green for their helpful comments; I. Higgins for technical assistance; C. Joyner for plant care.

