## Supplemental Figures and Tables for "Recruitment of *GRASSY TILLERS1 (GT1)* and *RAMOSA3 (RA3)* into distinct genetic networks in the evolution of grass morphology"

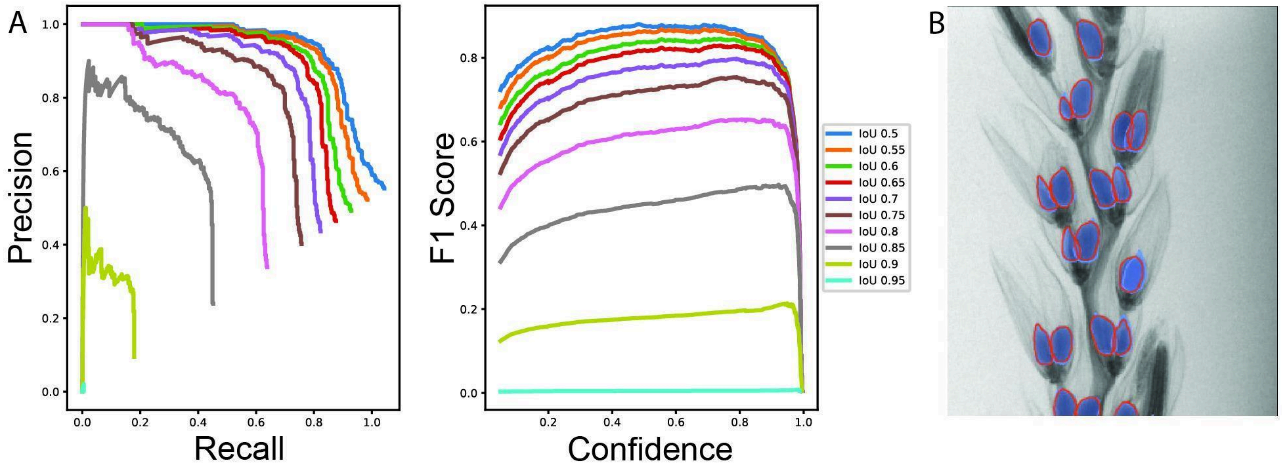


C

| IoU Mean / Median / Mode | Precision - AP50 | Precision - AP75 | Precision - mAP |
|---|---|---|---|
| .79, .83, .85 | .96 | .83 | .70 |
| **IoU Variance** | **Recall - AR50** | **Recall - AR75** | **Recall - mAR** |
| Q1: .73, Q2: .79, Q3: .89, IQR: .15 | .77 | .67 | .56 |

**Supplemental Figure 1**: **Model Performance Summary @ 0.82 Confidence Threshold.** A) Precision vs. Recall curves for final model performance on hold-out maize test dataset, split by IoU threshold. F1 Score vs. Confidence for final model performance on hold-out maize test dataset, split by IoU threshold. For this application we prioritized segmentation accuracy and precision over recall. The maximum F1 scores at high IoU thresholds occurred at or above 0.82, so we used 0.82 as the confidence threshold. B) Example detections (red outlines) and ground truth (blue shading) of a *gt1 ra3* maize plant at the 0.82 confidence threshold. C) Final model performance statistics at the 0.82 confidence threshold used for analysis. Segmentation precision and recall are reported at the .50 IoU threshold (AP50, AR50) and .75 IoU threshold (AP75, AR75). The mean average precision and recall (mAP, mAR) are calculated by taking the mean of the APs/ARs from the .50 to .95 IoU thresholds, in steps of .05.

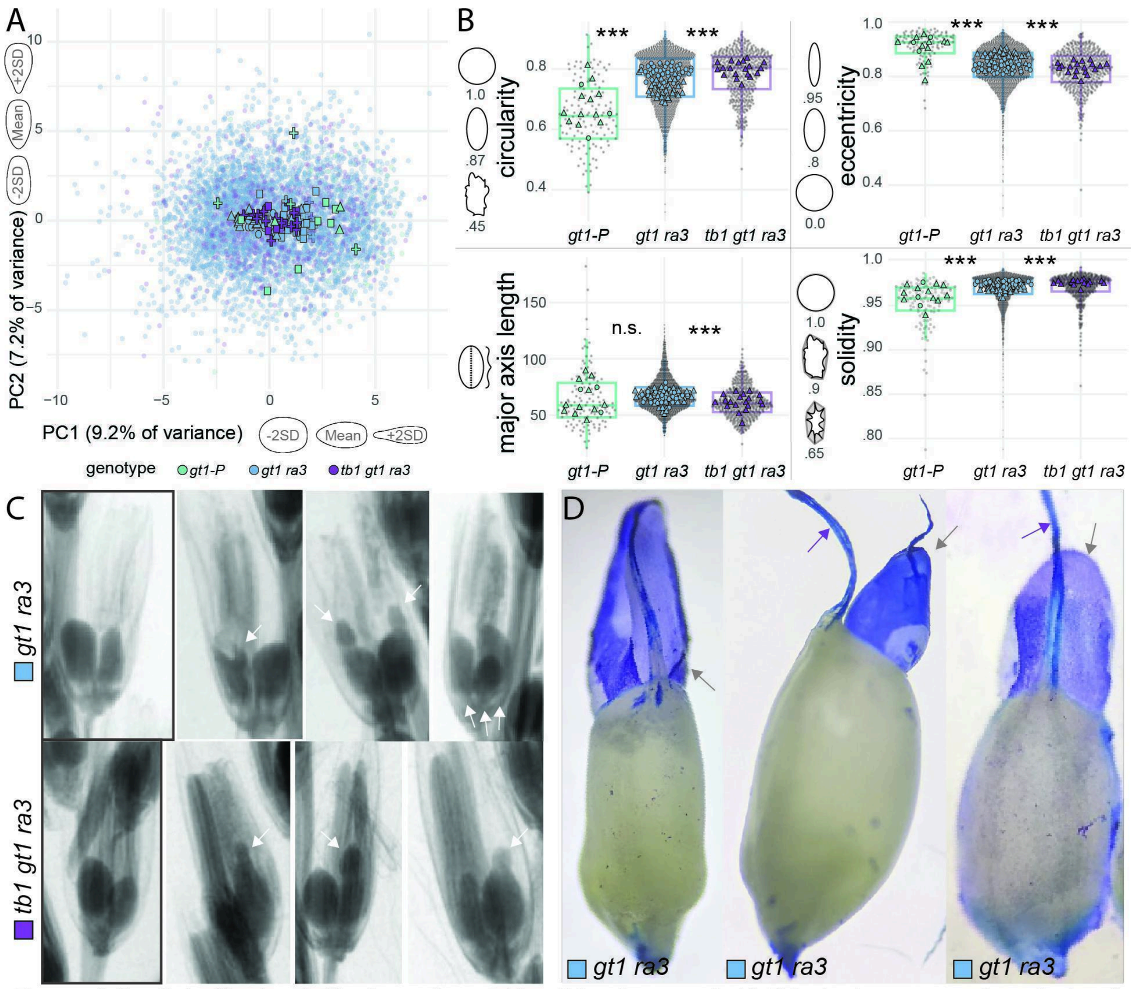


**Supplemental Figure 2. Pistil shape varies across genotypes**. A) Principal component analysis of elliptical Fourier pistil descriptors. Large shapes on the graph show the mean value of all pistils per tassel. B) Shape metric comparison with example shapes on the left. Genotype statistical comparisons were done with a mixed linear model: shape metric ~ genotype * block + (1 | plant) and estimated marginal means. *** = p-value < .05, n.s. = not significant. C) Examples of pistils from x-rays. Pistils with normal morphology are on the left. Pistils to the right are abnormal, with arrows indicating excess tissue. The last *gt1;ra3* spikelet shows three pistils instead of two. D) Dissected abnormal *gt1;ra3* pistils, stained with toluidine blue. Grey arrows indicate possible integument overgrowth, which is stained more darkly than the underlying nucellus. Purple arrows indicate silks.

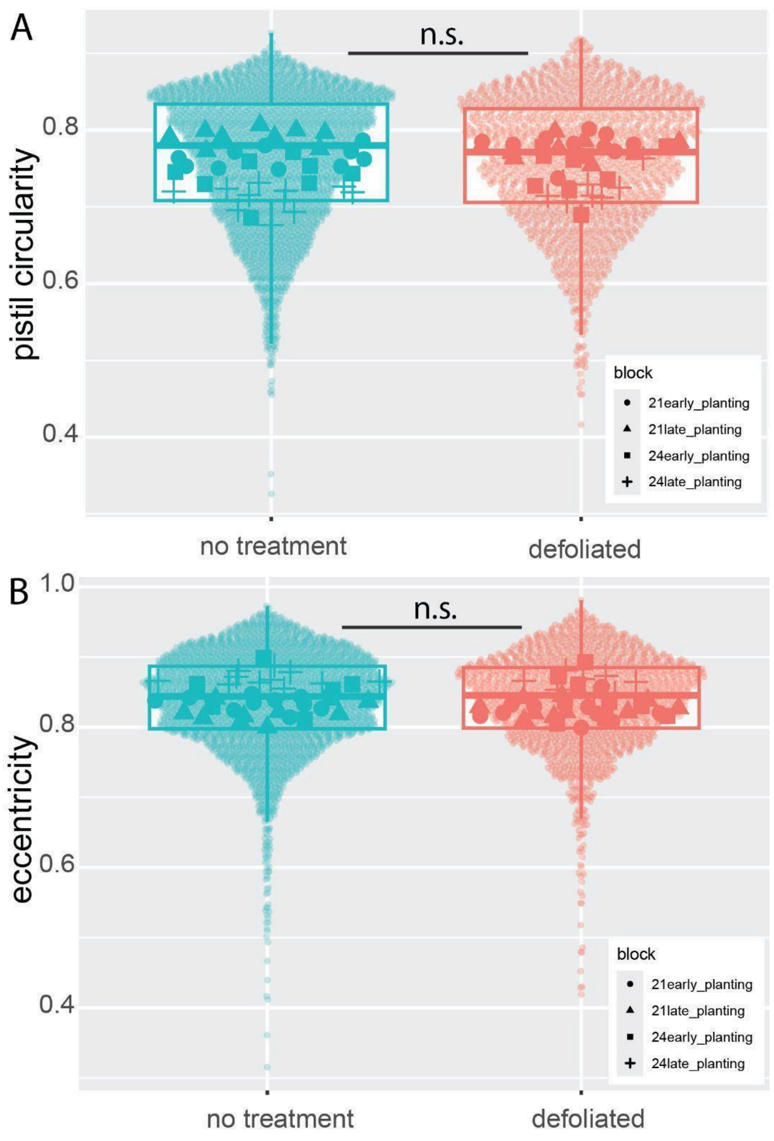


**Supplemental Figure 3. Defoliation of *gt1 ra3* plants does not affect pistil shape.** A) Pistil circularity compared between treatments. Large shapes show the average for each tassel. Significance test: Linear mixed model, circularity ~ treatment * block * (1 | plant). B) Pistil eccentricity compared between treatments. Significance test: Linear mixed model, eccentricity ~ treatment * block * (1 | plant)

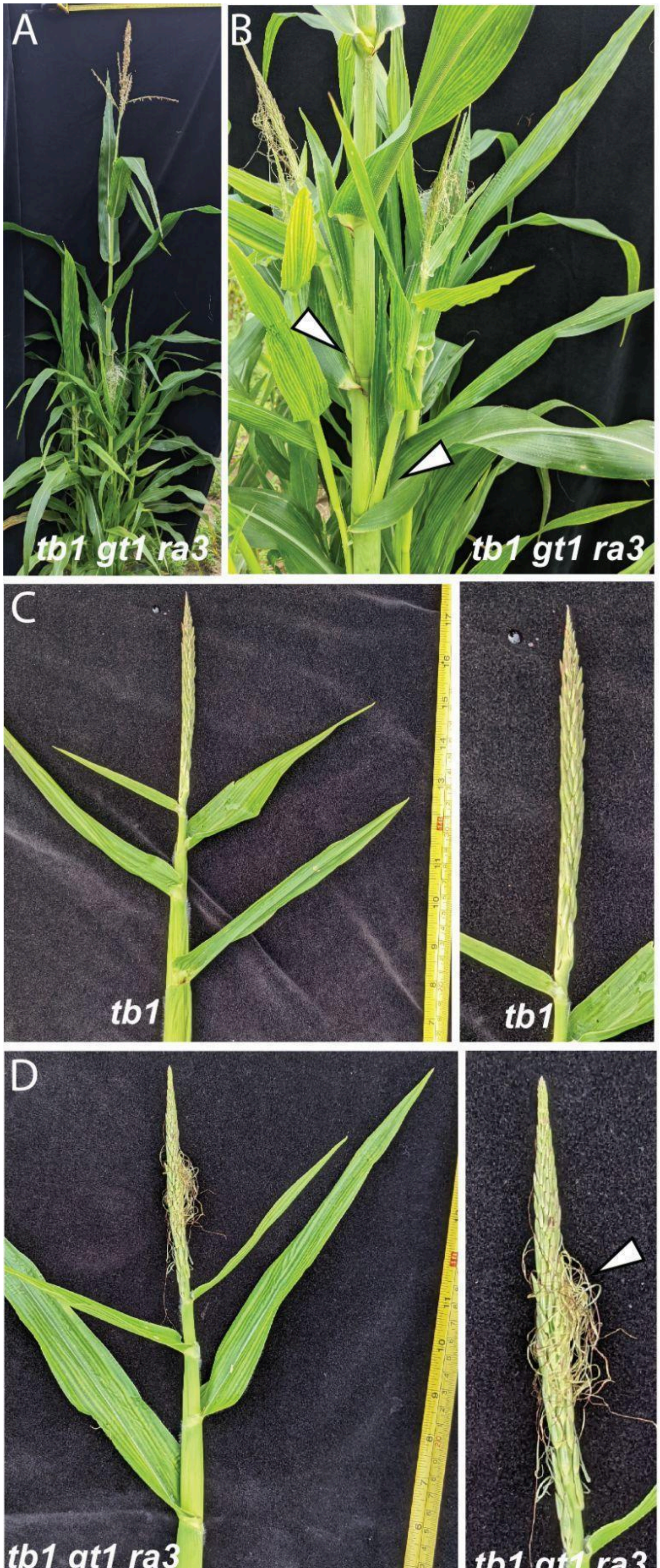

A
tb1 gt1 ra3
B
tb1 gt1 ra3
C
tb1
tb1
D
tb1 gt1 ra3
tb1 gt1 ra3

**Supplemental Figure 4. *tb1 gt1 ra3* phenotypes.** A) *tb1 gt1 ra3* whole plant, showing tillering. B) Close up of *tb1 gt1 ra3* middle section. White arrows show tiller-like structures in place of ears. C) A *tb1* tiller in place of an ear. D) A *tb1 gt1 ra3* tiller in place of an ear, arrow shows the silks of the derepressed carpels.

**Tables:**

| species | pistil size | genotype | stage | training images | validation images | test images |
|---|---|---|---|---|---|---|
| *Zea mays* | none | B73 (WT) | anthesis | 0 | 3 | 3 |
| *Zea mays* | none-small | *gt1-P* | anthesis | 33 | 8 | 7 |
| *Zea mays* | medium | *gt1 ra3* | anthesis | 64 | 12 | 11 |
| *Zea mays* | large | *tb1 gt1 ra3* (hybrid) | anthesis | 16 | 4 | 4 |
| *Setaria viridis* | none-small | A10.1 (WT) | 21 DAS | 29 | 7 | 6 |
| *Setaria viridis* | none-small | *svra3-1* | | | | |
| *Setaria viridis* | medium | A10.1 (WT) | 31 DAS | 32 | 6 | 6 |
| *Setaria viridis* | medium | *svra3-1* | | | | |
| *Setaria viridis* | large | A10.1 (WT) | 33 DAS | 30 | 6 | 6 |
| *Setaria viridis* | large | *svra3-1* | | | | |
| *Setaria viridis* | XL | A10.1 (WT) | 42 DAS | 27 | 6 | 6 |
| *Setaria viridis* | XL | *svra3-1* | | | | |

**Supplemental Table 1**: Overview of training, validation, and test datasets for model

| | **bbox AP50** | **bbox AP75** | **bbox mAP** | **segm AP50** | **segm AP75** | **segm mAP** |
|---|---|---|---|---|---|---|
| Fold 0 | 81.6 | 47.3 | 45.7 | 81.9 | 55.0 | 49.5 |
| Fold 1 | 83.1 | 51.4 | 47.8 | 82.8 | 55.5 | 50.2 |
| Fold 2 | 82.4 | 50.3 | 47.2 | 82.3 | 56.2 | 49.7 |
| Fold 3 | 83.1 | 52.0 | 48.1 | 83.3 | 55.6 | 50.1 |

| | | | | | | |
|---|---|---|---|---|---|---|
| Fold 4 | 81.0 | 48.0 | 46.3 | 81.0 | 52.7 | 48.0 |
| Mean ± SD | 82.2 ± 0.9 | 49.8 ± 2.1 | 47.0 ± 1.0 | 82.3 ± 0.9 | 55.0 ± 1.4 | 49.5 ± 0.9 |

**Supplemental Table 2:** 5-Fold stratified cross validation. bbox = bounding box, segm = segmentation.

| | **bbox AP50** | **bbox AP75** | **bbox mAP** | **segm AP50** | **segm AP75** | **segm mAP** |
|---|---|---|---|---|---|---|
| maize test set | 87.2 | 64.2 | 55.0 | 87.4 | 69.3 | 58.6 |
| setaria test set | 81.4 | 54.7 | 48.5 | 81.5 | 58.6 | 50.0 |

**Supplemental Table 3:** Model performance on hold-out test sets, no confidence threshold applied. bbox = bounding box, segm = segmentation.
